# Local DNA shape is a general principle of transcription factor binding specificity in *Arabidopsis thaliana*

**DOI:** 10.1101/2020.09.29.318923

**Authors:** Janik Sielemann, Donat Wulf, Romy Schmidt, Andrea Bräutigam

**Affiliations:** Computational Biology, Center for Biotechnology (CeBiTec), Bielefeld University, 33615 Bielefeld, Germany; Computational Biology, Faculty of Biology, Bielefeld University, 33615 Bielefeld, Germany; Graduate School DILS, Bielefeld Institute for Bioinformatics Infrastructure (BIBI), Bielefeld University, 33615 Bielefeld, Germany; Plant Biotechnology, Bielefeld University, 33615 Bielefeld, Germany

## Abstract

A genome encodes two types of information, the “what can be made” and the “when and where”. The “what” are mostly proteins which perform the majority of functions within living organisms and the “when and where” is the regulatory information that encodes when and where DNA is transcribed. Currently, it is possible to efficiently predict the majority of the protein content of a genome but nearly impossible to predict the transcriptional regulation. This regulation is based upon the interaction between transcription factors and genomic sequences at the site of binding motifs^1,2,3^. Information contained within the motif is necessary to predict transcription factor binding, however, it is not sufficient^4^, as experimentally verified binding sites are substantially scarcer than the corresponding binding motif. Thus, it remains challenging to derive regulational information from binding motifs. Here we show that a random forest machine learning approach, which incorporates the 3D-shape of DNA, enhances binding prediction for all 216 tested *Arabidopsis thaliana* transcription factors and improves the resolution of differential binding by transcription factor family members which share the same binding motif. Our results contribute to the understanding of protein-DNA recognition and demonstrate the extraction of binding site features beyond the binding sequence. We observed that those features were individually weighted for each transcription factor, even if they shared the same binding sequence. We show that the gained insights enable a more robust prediction of binding behavior regarding novel, not-in-genome motif sequences. Understanding transcription factor binding as a combination of motif sequence and motif shape brings us closer to predicting gene expression from promoter sequence.

## Main

Changes in gene expression during development and invoked by environmental perturbations are critical to organismal function and these changes are influenced by DNA-binding transcription factors. *Arabidopsis thaliana* encodes 1533 DNA-binding transcription factors^5^ many of which occur in protein families of a few to over a hundred members^6^. The DNA-binding transcription factors carry a domain which interacts directly with DNA as well as domains which mediate regulation. Gene expression of a particular gene is a complex read-out based on the presence of transcription factors and their spacing on the DNA, chromatin status, histone flags and presence of co-activators or repressors. Improving the understanding of those regulatory relations and pathways is necessary to tackle current agricultural challenges^7^. Hundreds of sequence motifs to which transcription factors bind have been characterized^3,8^, but currently it is impossible to look at a promoter and understand its regulatory syntax.

This is partially due to the fact that many members of a particular TF family bind the same motif. The *A. thaliana* genome contains 74 members of the WRKY transcription factor family^9^. These WRKY transcription factor family proteins are implicated in diverse processes from trichome and seed development to roles in biotic and abiotic stresses^10^. Despite their diverse roles, all WRKYs analyzed bind to a consensus motif, the W-box, which is characterized by the TTGAC pentamer followed by C or T^11^. Transcription factors of the 133 member bHLH family bind DNA via basic amino acids at the N-terminal end of the bHLH domain and bind a variation of the motif CANNTG, frequently the so-called G-box CACGTG^12^. This G-box motif is also bound by many bZIP transcription factors whose core motif is ACGT, the central nucleotides of the G-box CACGTG^13,14^. ChIP-seq data clearly indicates that only a subset of potential binding sites are indeed occupied at any given time in a particular tissue^15–17^. To explain why transcription factors binding the same sequence yield different regulatory output, additional variables such as other transcription factors as binding partners, chromatin status, histone modifications, DNA methylation and posttranslational modifications of transcription factors have been considered. In contrast, the functional divergence has been hypothesized to be dependent on varying binding affinities to the same core sequence^4^.

In DNA affinity purification sequencing (DAP-seq) experiments the DNA bound to an *in vitro* produced transcription factor is sequenced. It has also revealed extensive family-type motifs for many transcription factors (http://neomorph.salk.edu/dap_web/pages/browse_table_aj.php, ^3^). For motif detection in DAP-seq or ChIP-seq, the DNA sequences bound by the TF are mined by motif search algorithms such as MEME or MEME-chip which identify overrepresented motifs among the sequences^18^. For many TFs, biochemical experiments such as EMSAs have confirmed that the motif is necessary for binding^19–21^. The comparison of the direct results of DAP-seq, i.e. the peaks in which reads can be aligned with the genome with the occurrence of the motif predicted using the same reads indicates that during motif prediction information is lost (Figure 1A, Supplemental Figure 1). AmpDAP-seq uses a single *in-vitro* produced protein that binds amplified DNA devoid of methylation marks, histones and other proteins. Hence the bound DNA sequence contains information which is lost during motif prediction. This raises the question which information is contained in DNA sequence besides the code of A, T, C and G.

**Figure 1:**
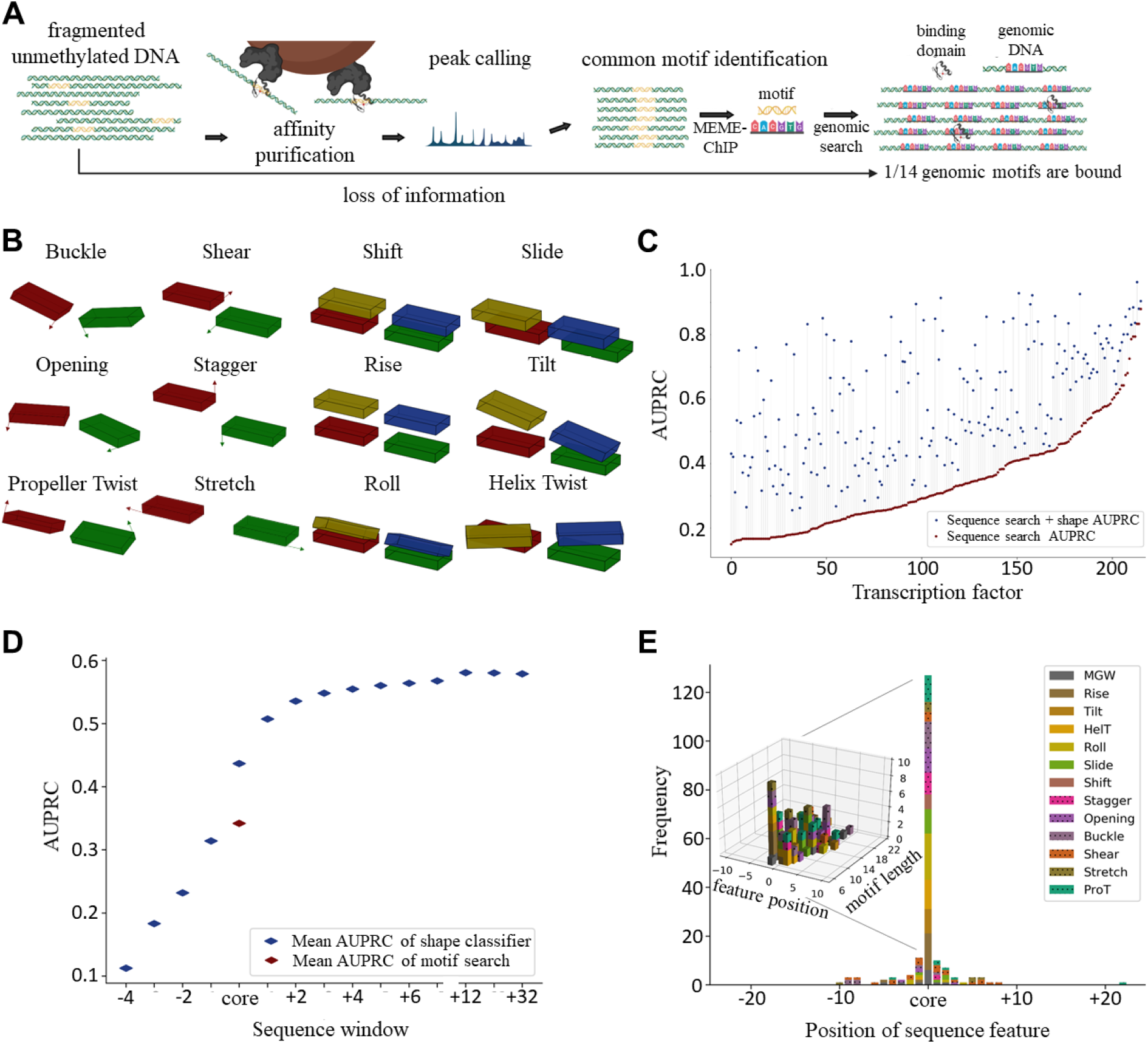
Overview of workflow and performance of shape based binding site identification. A) Broad overview of the experimental procedure for the identification of TF binding sites. The DAP-Seq experiment was performed by O’Malley et al. 2016. Identified binding motif occurrence is on average 14-fold higher than the number of verified binding events. B) The different DNA shape features which were considered to analyse TF specificity. A query table was used for shape calculation^22^. C) Performance of the random forest classifier using the border widths with the highest area under the precision recall curve (AUPRC) for each transcription factor. D) AUPRC for differing sequence widths. The width was increased upstream and downstream of the core motif sequence, respectively. E) Frequency of features in the top 5 most important shape features for each transcription factor.

DNA is a very constrained molecule since its phosphate sugar backbone runs antiparallel while its bases are paired and arranged in rungs on a helical ladder. Since the discovery of the genetic code for proteins, we have been trained to see DNA essentially as code. However, despite the constraints, the exact position of each base pair and each base in a pair is influenced by its surrounding bases. The pairs can be tilted, shifted, slid, rolled, risen and twisted relative to each other (Figure 1B,^22,23^). The bases in a pair can be buckled, sheared, stretched, twisted, opened and staggered (Figure 1B,^22^). The width of the minor groove is also influenced by the surrounding bases^24^. This DNA shape has been demonstrated to influence protein-DNA binding, for instance of the *Drosophila* Scr Hox Protein^24,25^ and the *S. cerevesia* bHLH proteins Cbf1 and Tye7^26^. We hypothesized, that models built on DNA shape within and surrounding the binding motif recover the information lost during motif detection and improve prediction for transcription factor binding in *A. thaliana*.

DNA shape calculations have been established from X-ray crystallography, Monte Carlo simulation and molecular dynamic simulation^22^. DNA shape is encoded in 13 features represented by numeric vectors. To test the importance of DNA shape, a random forest decision tree (RF) based machine learning approach was chosen to enable reliable feature extraction^27^. To test whether DNA shape encodes additional information, we split the motif occurrences into a training and a test-set^28^ using the peak height of a DAP-seq experiment as the numeric for which the RF algorithm was trained. To ensure consistent 3D structure learning, the sequences were reverse complemented if the binding sequence was located on the minus strand. Initially, a random forest classifier was trained using the binding/non-binding cut-offs established previously^3^. We observed improved binding predictions for all tested transcription factors (Supplemental Figure 2), but quantitative information was lost. In a second step, a regressor was trained on the raw binding data using the peak height in ampDAP-seq as a proxy for binding affinity. The performance was similar (Supplemental Figure 3). To improve the use of peak height as a proxy for binding affinity, the binding data was filtered for single motif occurrences and machine learning was repeated. 216 transcription factors were tested and in each case the shape based predictor outperformed the motif search based on the area under the precision recall curve (AUPRC). AUPRC improved between 2.8% and 362.7%, with an average of 93.2% (Figure 1C). 33 TFs reach AUPRC of more than 0.8 indicating that the motif plus shape information suffices for prediction (Figure 1C). 101 TFs show medium AUPRC with between 0.5 to 0.8 AUPRC. The remaining 82 TFs improve their AUPRC compared to motif alone but do not exceed an AUPRC of 0.5 (Figure 1C). Prediction of binding improved for all transcription factor families, however, some families increased in prediction precision more than other (Supplemental Figure 4, Supplemental Figure 5). The test-sets clearly indicate that DNA shape contributes to TF binding specificity when bound vs non-bound sequences are analyzed (Figure 1C). An additional comparison between ampDAP-seq and ChIP-seq data was performed for 5 transcription factors, for which data of both experimental procedures were available (Supplemental Figure 6). We observed that ampDAP-seq outperformed ChIP-seq for each TF with an average of 98.6% higher AUPRC. To test the contribution of shapes surrounding the motif, the amount of shape information given to the regressor was varied and the training repeated. The major contribution of shape information was localized to the core motif plus two bases on each side of the motif (Figure 1D). These adjacent bases influence the shape of the bases and base pairs in the core^22^. Beyond the core motif shape, the information gain quickly leveled (Figure 1D, Supplemental Figure 7). The models generated by the shape-based regressor yield information over which shapes are important to the binding for each of 216 TFs tested (Supplemental Table 1). To test if any shape, shape type (intra-basepair vs. inter-basepair) or any position contribute a larger amount of information to the binding, the top five features were extracted for each transcription factor. The 3D configuration of bases within the core binding sequence occupied 68% of the top five feature positions (Figure 1E) as expected (Figure 1D). Intra-basepair shapes contributed 39% and inter-basepair shapes contributed 61% (Figure 1E) of the top five shapes within the motif. Outside of the motif, the proportions reversed since intra-basepair shapes contributed 72% and inter-basepair shapes contributed 28% (Figure 1E). Far outside the motif, the shear feature was overrepresented among the top five features (Figure 1E). More precise prediction of TF binding requires shape information and the transcription factors depend on a wide variety of shape information.

If DNA shape predicts binding better than motif alone and shape information used by TFs is varied, the prediction algorithm should be able to distinguish binding between two transcription factors which are predicted to bind the same motif sequence. To test the hypothesis, the models for transcription factor pairs with the same binding motifs were analyzed. The ERF/AP2 transcription factors CBF4 (AT5G51990) and ERF036 (AT3G16280) both bind the GTCGGT/C motif which occurs 31,155 times in the *A. thaliana* genome. According to ampDAP-seq they have 9,910 binding sequences in common (Figure 2A). ERF036 binds 2,581 sequences which are not bound by CBF4, and CBF4 binds 6,996 sequences not bound by ERF036. To test whether the shape indeed supplies specificity, binding vs non-binding was predicted by the models (Figure 2B).

**Figure 2:**
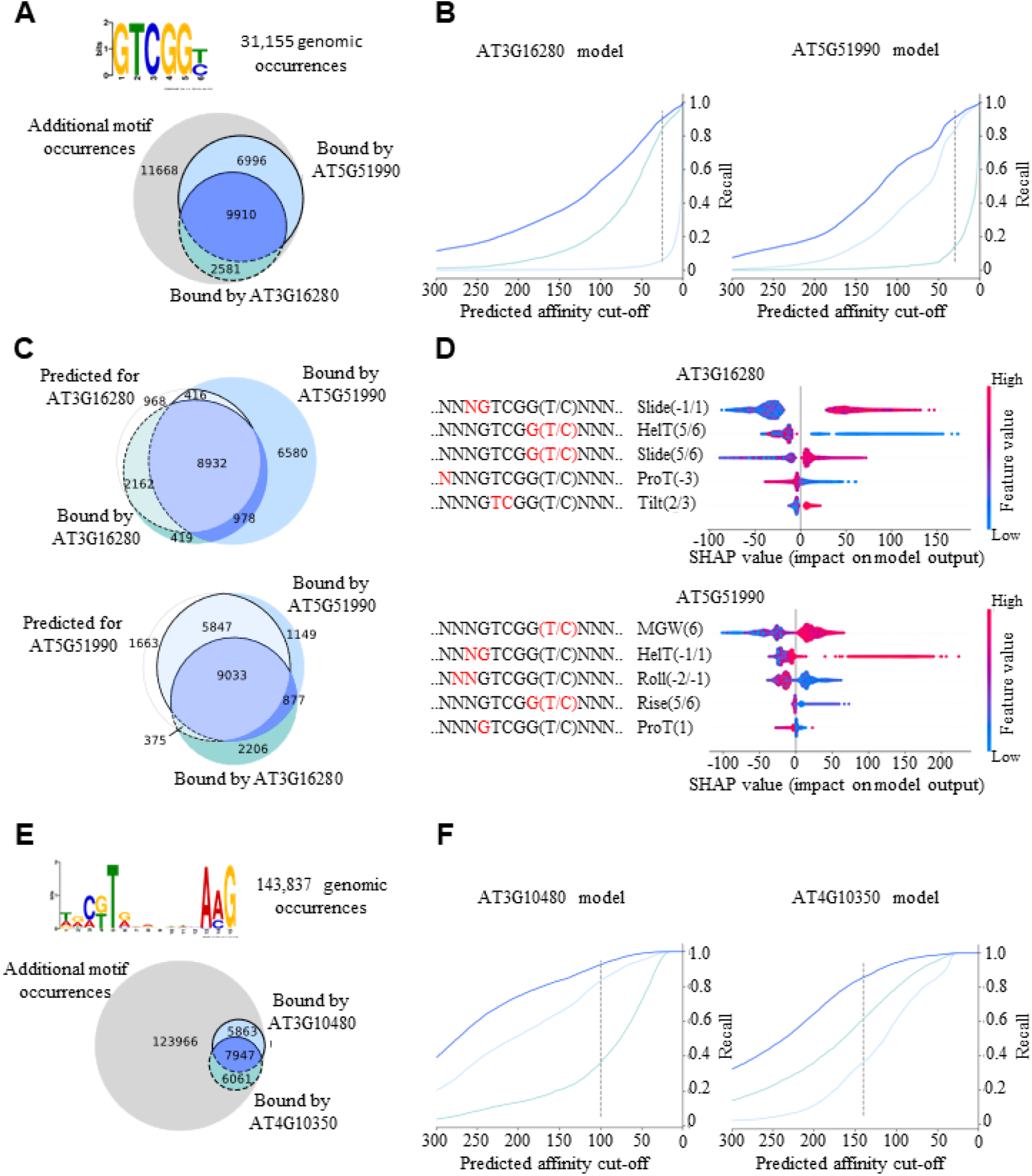
Differentiation of binding specificity of intra-familiar proteins with the same binding motif. A, E) Occurrence of the GTCGG(T/C) and C(G/T)TNNNNNNNAAG binding motifs in the *A. thaliana* genome sequence and the experimentally validated binding sequences of the AP2/EREBP TFs AT5G51990 and AT3G16280 and NAC TFs ANAC050 (AT3G10480) and BRN2 (AT4G10350). B, F) Performance of the random forest regressor trained on the genomic 3D shape. Each line represents the ratio of correctly predicted binding sites regarding all validated binding sites for different affinity prediction cut-offs. The dark blue line corresponds to binding sequences which are bound by both transcription factors and the light blue lines correspond to the uniquely bound binding sequences. C) The Venn diagrams show the sequence distributions according to the cut-off represented by the dashed line, respectively. Fields with light colours show the overlap of predicted and validated binding sequences. Dark coloured fields show the quantity of sequences, which were not predicted as bound by the model regarding the shown cut-off. (D) Influence of different local shape features on the prediction of the regressor model. The most influential features are at the top. Each row represents one shape feature at a single position within the sequence.

The shape information of the 31,155 genomic sequences, allows the regressor models to distinguish binding events between the two transcription factors (Figure 2B-C), even though the core sequence is the same and the majority of sequences are bound by both transcription factors. Each Venn diagram in Figure 2C shows the distribution of binding sites applying the cutoff represented by the dashed line. For ERF036 2,162 out of the 2,581 uniquely bound sequences were correctly identified as binding sequences, whereas only 416 out of the 6,996 sequences bound uniquely by CBF4 were wrongly predicted as binding sequences (Figure 2C). In total, the number of false positive binding sequences dropped from 18,664 (11,668+6,996) to 1,384 (968+416), which is an improvement of 93%. Likewise, for CBF4 84% of uniquely bound sequences were correctly predicted and only 15% of sequences bound uniquely by ERF036 are predicted as false positives (Figure 2C). Here, the total improvement regarding false positives amounts to 86% (2,038/14,249). To identify the features which contribute specificity to each TF, we extracted feature importances using ‘shapley additive explanations’ (SHAP)^27^ (Figure 2D). The outputs of the regressor models are influenced by different features. For ERF036 the slide at position −1 relative to the motif and the helix twist at position 5 in the motif is most influential, whereas for CBF4 the minor groove width at position 6 and the helix twist at position −1 contribute most to the decision of the random forest model. This observation underlines that the transcription factors, even though binding to the same core motif, are dependent on different peculiarities regarding the shape of the DNA (Figure 2). These results are not family specific since transcription factors of the NAC family binding to the C(G/T)TNNNNNNNAAG motif (Figure 2E-F), transcription factors of the WRKY family binding to TTGAC(T/C) motif, transcription factors of the bZIP family binding to ACGTCA motif and transcription factors of the C2H2 family binding to TTGCTNT motif show similar results (Supplemental Figure 8-10). In summary, the features defined by the shape-based regressor are able to explain differential binding of two transcription factors binding to the same sequence motif.

When used in a train-test scenario, the models generated by machine learning improve binding site prediction (Figure 1) and distinguish binding events for TFs binding the same motif (Figure 2). To test if the models are able to produce novel information they were used to predict TF binding to sequences not present in the *A. thaliana* genome. For the HY5 (AT5G11260) transcription factor of the bZIP family with the core motif ACGT, six DNA sequences with high (>150 peak height units) and low (<15 peak height units) regressor predictions were generated. For this purpose, 100,000 sequences not present in the genome of *A. thaliana* consisting of 18 bases with ACGT as core sequence were randomly created and the regressor model was applied. Similarly, six DNA sequences were generated for the transcription factor ANAC050 (AT3G10480). The binding affinity was experimentally tested by performing an electrophoretic mobility shift assay (EMSA) (Figure 3A-B, Supplemental Figure 11, Supplemental Figure 12). Without any competitor, a shifted band is visible compared to the negative control (Figure 3A-B). In the comparative competition experiment with HY5, using shapes with low regressor values, all labelled bands are still visible (Figure 3A). Those shapes are not able to out-compete the labelled sequence and are thus apparently not bound with high affinity by HY5. For the shapes with high regressor values, two out of three do not show any labelled band and are therefore bound by HY5 with sufficient affinity to out-compete the labelled sequence. Predictions are thus in the predicted error probability (PRAUC=0.72; 5/6=0.83). For ANAC050 the EMSA shows similar results with five out of six predictions being correct (Figure 3B). In total, we observed that 10 out of 12 predictions were experimentally validated for both transcription factors. Given that the AUPRCs for both proteins yielded 0.72 and 0.78 (Figure 3C-D), the validation of binding and non-binding events occurs within the expected error rate. To illustrate the subtle relevant factors, schematic models of the DNA sequences were plotted. For HY5, the schematic model of base and base pair shape shows visually obvious differences on the buckle at position +3 and the shear of position −1 between bound and not bound sequences (Figure 3C). Additionally, important positions for binding extracted with SHAP are the helix twist at positions 5 and +1 and the opening at position −1. (Figure 3C, Supplemental Figure 13). For the ANAC050 protein, the most obvious difference between the bound and not bound sequence is that the bound sequence is overall more stretched out. The main reason for this observation is that the average roll for the bound sequence is approximately −0.88°, whereas the bases of the sequence which is not sufficiently bound are rolled on average by approximately −1.77° (Figure 3D).The EMSA confirmed the predictive capability of the model constructed by machine learning.

**Figure 3:**
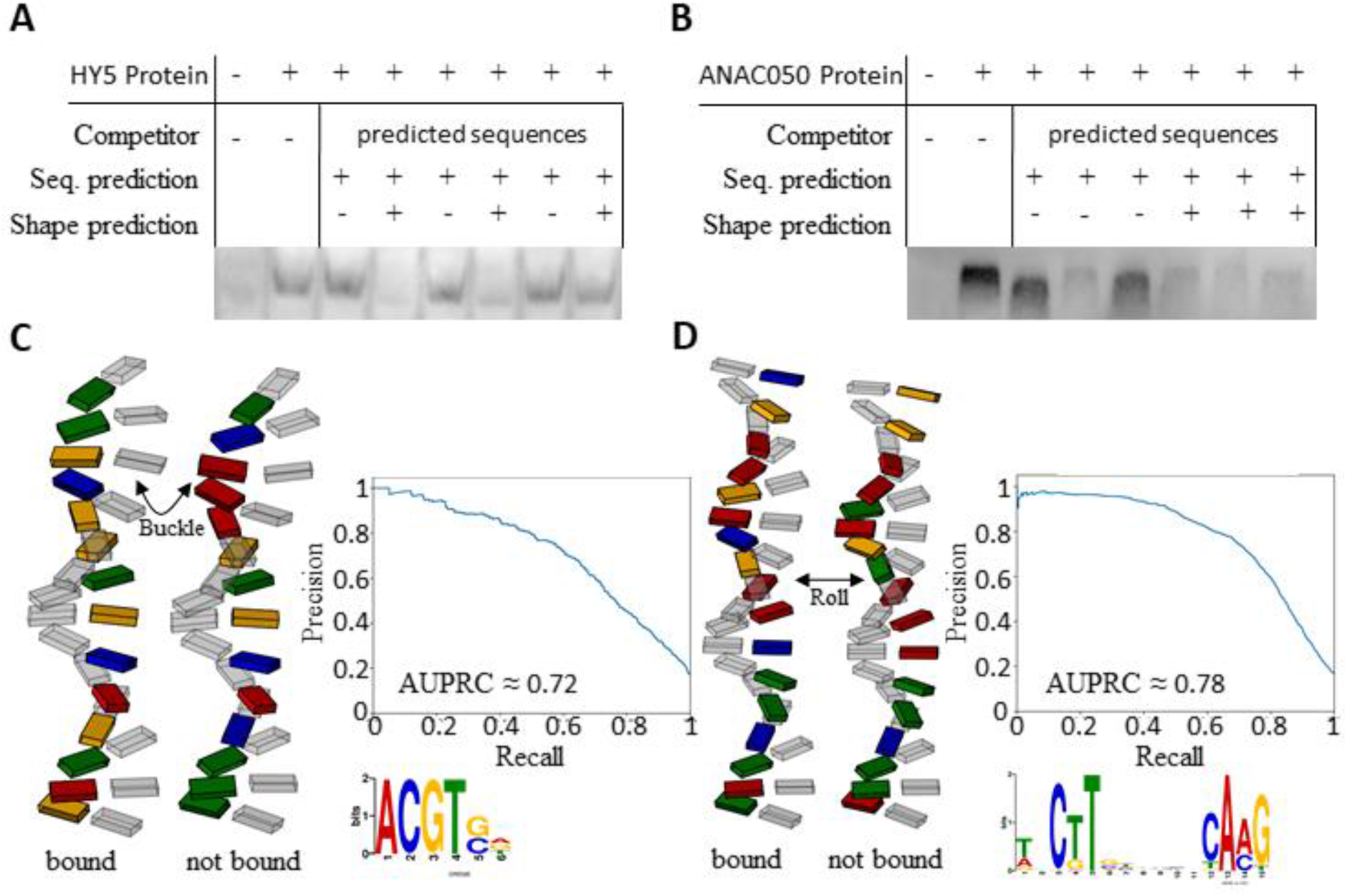
Experimental validation of shape based prediction for HY5 binding sequences. A,B) Competition EMSA for sequences containing the sequence motif for each respective transcription factor with high and low binding affinity predictions based on their 3D structure.. C,D) Illustration of the 3D structure of the corresponding sequences. The DNA backbone is not shown, as it is not yet possible to reliably calculate the spatial arrangement of the backbone. Additionally, the precision-recall curve of the random forest models for the respective transcription factors are shown. Precision and recall are based on the ampDAP *in vitro* verified binding sequences.

Our results show that the binding behaviour of transcription factors depends on the 3D formation of its binding site, where different transcription factors favour different formations even within the same protein family. In contrast to ChIP-seq data, ampDAP-seq data, which uses naked genomic DNA^3^, allows a more precise identification of 3D feature importances for each transcription factor individually. Large experimental efforts have been designed to precisely assign binding sites to transcription factors^29^ and to use this knowledge to describe transcriptional regulation^30^. The experiments (Figures 1–3) show that a combination of motif sequence and motif shape enables improved prediction of transcription factor binding on genomic sequence. The models generate a catalogue of potential binding sites in a genome and their predicted affinity. This information forms a base on which additional information layers (i.e. spacing of binding sites^31^, chromatin openness^32^, histone flags^32^, and quantity of transcription factors and their interactors) can be stacked to enable prediction of gene expression. In synthetic biology, binding events for heterologously expressed transcription factors can be predicted more precisely and rationally designed promoter sequences are one step closer.

In the future, it will be critical to study evolutionary trajectories of transcriptional regulation to determine changes to binding sites present in genomes and changes to shape preferences of transcription factors. Precise understanding of transcription factor binding will allow us to build predictive regulatory networks and hence enable us to understand agriculturally important complex traits such as differential responses to heat, drought and pathogens and control of yield.

## Methods

The ampDAP-seq peak calling data were obtained from the Plant Cistrome Database (neomorph.salk.edu/dap_web/pages/index.php)^3^. Only datasets with a ‘fraction of reads in peaks’ (FRiP) value > 5% were considered for further analyses. All peak sequences were extracted from the *A. thaliana* reference genome sequence (TAIR10), obtained from ftp://ftp.arabidopsis.org/. The peak sequences were then used as input for the MEME-ChIP tool^33^ to detect binding motifs. The motif with the lowest e-value was chosen as core motif for each transcription factor. Peaks, which appeared in more than one third of all data sets, are considered as artifacts and were discarded.

To determine motif frequency in the genome, core motif occurrences were searched within the *A. thaliana* genome sequence using FIMO^34^. All motifs located within 80 base pairs of a peak summit were considered as experimental validated binding events. Multiple motif occurrences within this defined peak area were classified as homodimer binding sites to enable a more precise signal value interpretation. The calculation of the DNA shape was performed using a query table extracted from the Bioconductor package DNAshapeR^22,35^.

The random forest classifier as well as the random forest regressor models were generated and trained using the python module scikit-learn^28^. Hyperparameter grid search and 5-fold cross validation were performed to generate each model. A detailed explanation of the data preprocessing and model generation is provided in the Supplemental Methods. Code is available from GitHub (https://github.com/janiksielemann/shape-based-TF-binding-prediction).

The HY5 (AT5G11260) and ANAC50 (AT3G10480) coding sequence was cloned with Gibson assembly in pFN19A HaloTag® T7 SP6 Flexi® Vector (Promega, Madison, Wisconsin, United States) in an N-terminal fusion with the Halo-tag. Plasmid-DNA was isolated with the ZymoPURE Plasmid Midiprep (ZymoGenetics, Seattle, Washington, United States The HY5 protein was expressed with TnT® SP6 High-Yield Wheat Germ Protein Expression System (Promega, Madison, Wisconsin, United States) using 2 μg Plasmid DNA per 50 μL expression reaction. The ANAC50 protein was purified with the HaloTag^®^ Protein Purification System (Promega, Madison, Wisconsin, United States) using 20 μL Expression reaction for each EMSA reaction.. Expression was validated by Halo-tag detection (Supplemental Figure 14). Double stranded DNA sequences (20 μM) were generated by annealing synthesized DNA (98-21 °C, 9 hours) and diluted to 0.25 μM. The binding reaction was incubated for 2 h at 21°C. A 5% native polyacrylamide gel containing 0.5 TBE and 2.5% glycerol was prerun for 30 minutes. The samples were loaded with 1 μL Orange loading dye (Thermo Fisher Scientific, Waltham, Massachusetts, United States) and the gel (10 × 7.5 cm) was run at 80 V until the OrangeG front was 1 cm before the end of the gel. The gel was blotted on a positively charged nylon membrane (Hybond™, GE Healthcare, Chicago, Illinois, United States) at fixed current of 0.8 mA/cm^2^ for 90 minutes. The DNA was fixed by UV for 10 minutes. Biotin labeled DNA was detected with 1:5000 solution of an anti-biotin HPR-conjugated antibody (BioLegend, San Diego, California, United States) in TBST with 5% BSA. Detection was performed using Pierce™ ECL Western Blotting Substrate (Thermo Fisher Scientific, Waltham, Massachusetts, United States) as described by the manufacturer and the imaging system Fusion Fx7 (Vilber, Collégien, France).

## Acknowledgements

We thank the Bioinformatic Resource Facility team at the Center for Biotechnology (Bielefeld University) for technical support.

## Author contributions

JS designed and carried out the computational experiments including programming, interpreted the data, and co-wrote the manuscript, DW designed and carried out the wet lab experiments, interpreted data, and edited the manuscript, RS assisted with the wet lab experiments, interpreted data, and edited the manuscript, AB conceived the initial idea and the study, interpreted data, and co-wrote the manuscript

**No competing interests are declared.**

**Supplementary Information** is available for this paper

