## SupplementalFigure10 for "Local DNA shape is a general principle of transcription factor binding specificity in *Arabidopsis thaliana*"

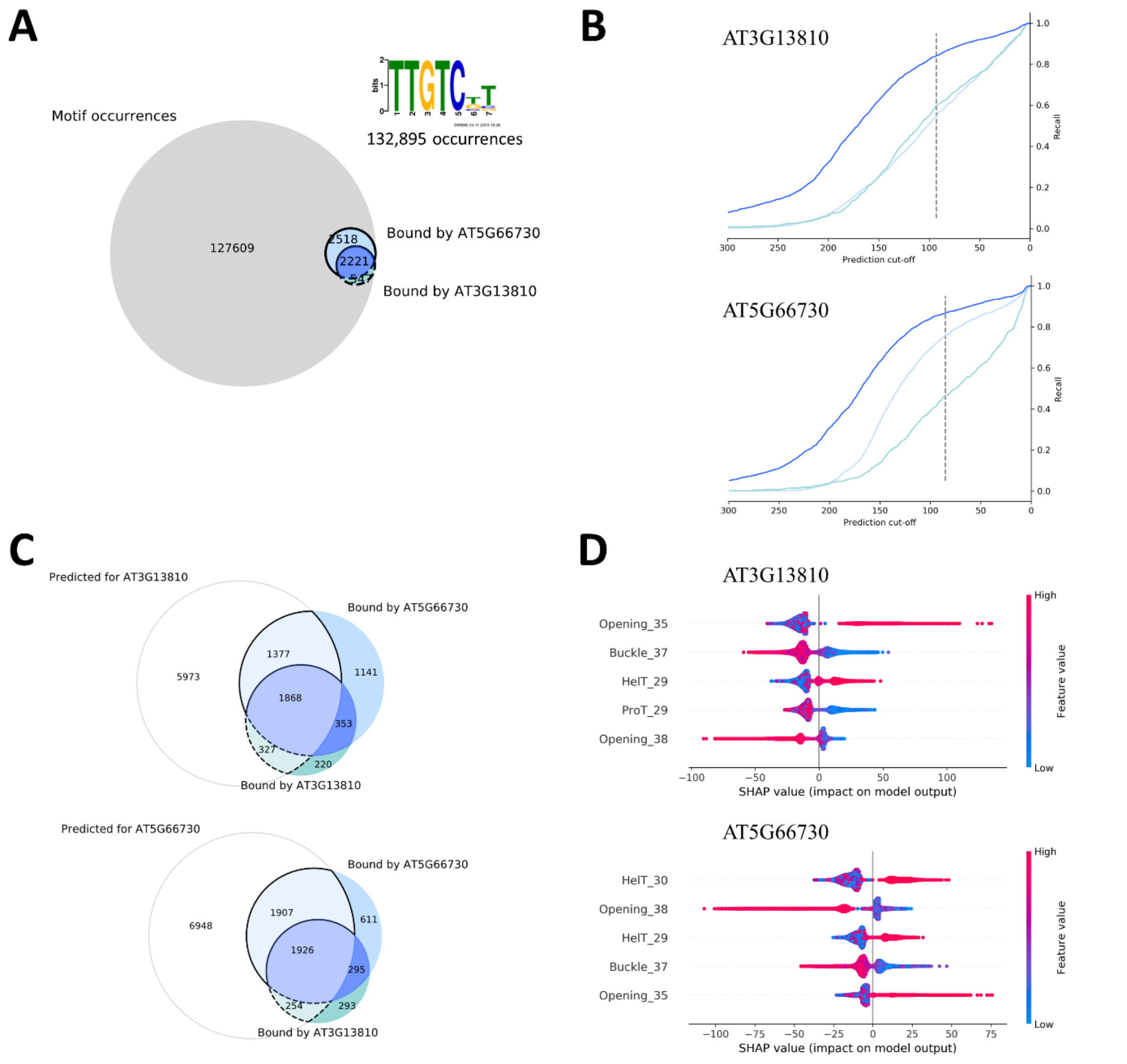


**Supplemental Figure 10: Differentiation of binding specificity of two C2H2 transcription factors with the same binding motif.**

A) Occurrence of the TTGTC(T/G)T binding motif in the *A. thaliana* genome sequence and the experimentally validated binding sequences of the C2H2 TFs AT5G66730 and AT3G13810. B) Performance of the random forest regressor trained on the genomic 3D shape. C) The Venn diagrams show the sequence distributions according to the cut-off represented by the dashed line, respectively. Fields with light colours show the overlap of predicted and validated binding sequences. Dark coloured fields show the quantity of sequences, which were not predicted as bound by the model regarding the shown cut-off. D) Impact of different local shape features on the prediction of the regressor model. The most influential features are at the top. Each row represents one shape feature at a single position within the sequence. The start of the core motif is at position 30.
