## SupplementalFigure11 for "Local DNA shape is a general principle of transcription factor binding specificity in *Arabidopsis thaliana*"

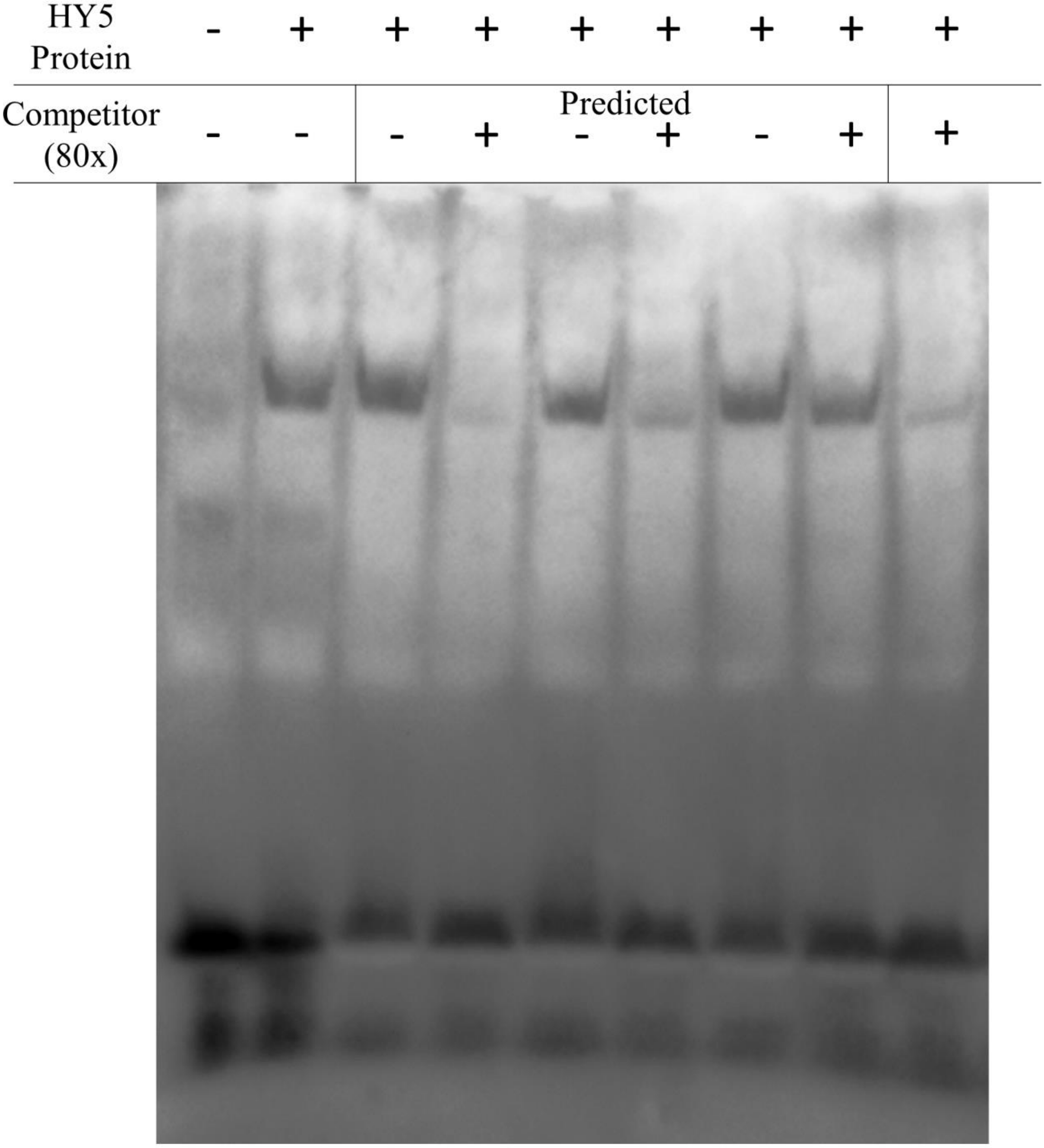


**Supplemental Figure 11: Competition EMSA with the *A. thaliana* transcription factor HY5.**

For experimental validation of the regressor predictions, DNA sequence with a high and low regressor prediction containing the same sequence motif not present in the genome of *A. thaliana* were generated. One sequence with a high prediction was labeled with biotin to detect DNA binding of HY5 by performing an EMSA. To compare binding affinities three sequences with high and low regressor predictions were used as competitors for the labeled sequence containing the same sequence motif. A shifted band is visible for all samples with low binding affinity prediction, one sample with high affinity prediction and the control without competitor sequence. Less visible shifted bands are observed in two samples with high predicted affinity and the control with the same competitor sequence.
