## SupplementalFigure12 for "Local DNA shape is a general principle of transcription factor binding specificity in *Arabidopsis thaliana*"

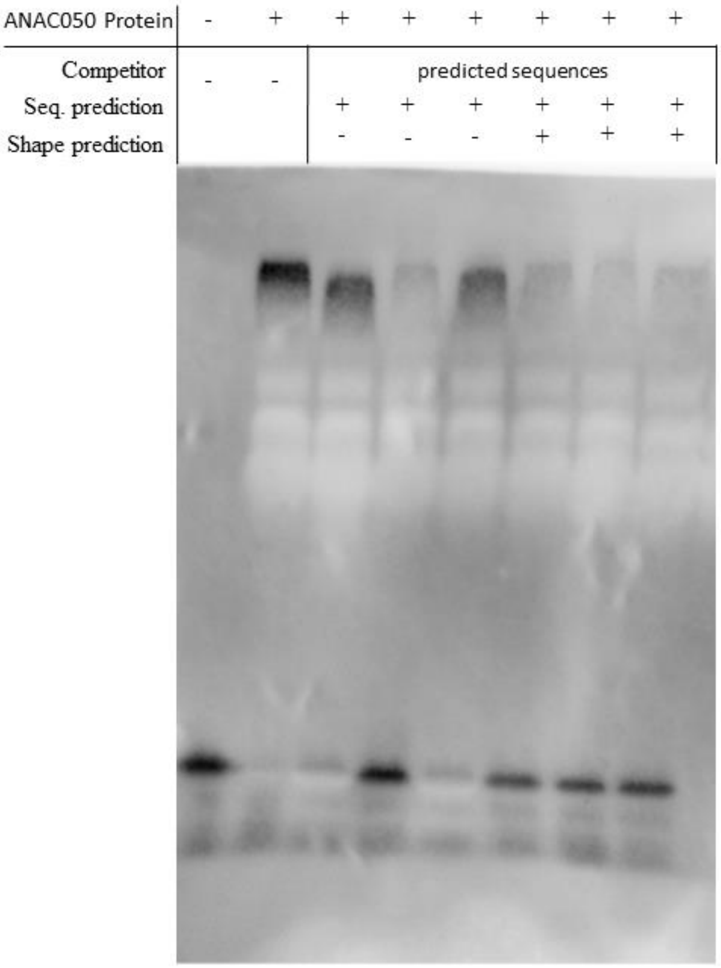


**Supplemental Figure 12: Competition EMSA with the *A. thaliana* transcription factor ANAC050.**

For experimental validation of the regressor predictions, DNA sequences not present in the genome of *A. thaliana* were generated. All sequences contain the extracted sequence motif for ANAC050. One sequence with a high regressor prediction was labeled with biotin to detect DNA binding of ANAC050 by performing an EMSA. To compare binding affinities three sequences with high and low regressor predictions were used as competitors for the labeled sequence. A shifted band is visible for two out of three samples with low binding affinity prediction and the control without competitor sequence. Less visible shifted bands are observed in all samples with high predicted affinity.
