## SupplementalFigure13 for "Local DNA shape is a general principle of transcription factor binding specificity in *Arabidopsis thaliana*"

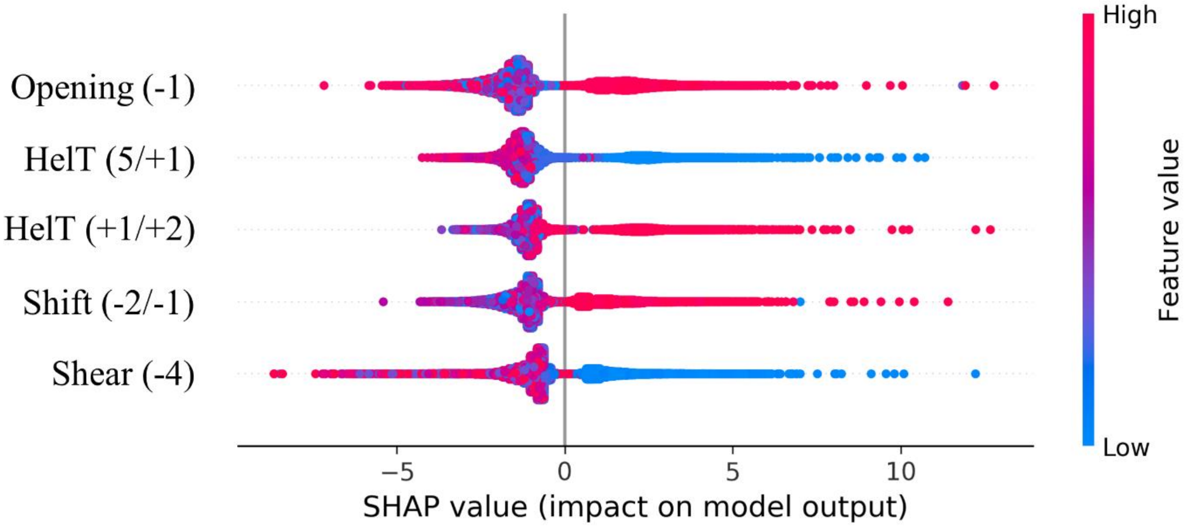


**Supplemental Figure 13: Top 5 most important shape features for HY5 binding.**

The SHAP Python package[1] was used to extract the most important features of the random forest model trained on experimentally validated HY5 binding sequences. The most important shape feature is the Opening at the -1 position, which is one base upstream of the motif sequence.
