## SupplementalFigure14 for "Local DNA shape is a general principle of transcription factor binding specificity in *Arabidopsis thaliana*"

The HY5 (AT5G11260) and ANAC050 (AT3G10480) coding sequences were cloned with Gibson assembly in the pFN19A HaloTag® T7 SP6 Flexi® Vector (Promega, Madison, Wisconsin, United States) in an N-terminal fusion in-frame with the Halo-tag. Plasmid-DNA was isolated with the ZymoPURE Plasmid Midiprep Kit (ZymoGenetics, Seattle, Washington, United States). The HY5 protein was expressed with the TnT® SP6 High-Yield Wheat Germ Protein Expression System (Promega, Madison, Wisconsin, United States) using 2 µg plasmid DNA per 50 µL expression reaction. The ANAC050 protein was purified with the HaloTag® Protein Purification System (Promega, Madison, Wisconsin, United States) using 20 µL expression reaction for each EMSA reaction. DNA sequences (20 µM) were annealed (98-21 °C, 9 hours) and diluted to 0.25 µM. Binding reaction was incubated for 2 h at room temperature. A 5 % native polyacrylamide gel containing 0.5 TBE and 2.5 % glycerol was prerun for 30 minutes. The samples were loaded with 1 µL Orange loading dye (Thermo Fisher Scientific, Waltham, Massachusetts, United States) and the gel (10 x 7.5 cm) was run at 80 V until the OrangeG band was 1 cm before the end of the gel. The gel was blotted on a positively charged nylon membrane (Hybond™, GE Healthcare, Chicago, Illinois, United States) at fixed current of 0.8 mA/cm$^2$ for 90 minutes. The DNA was fixated by UV with a transluminator for 10 minutes. Biotin labeled DNA was detected with a 1:5000 solution of an anti-biotin hpr conjugated antibody (BioLegend, San Diego, California, United States) in TBST with 5 % BSA. Detection was performed using Pierce™ ECL Western Blotting Substrate (Thermo Fisher Scientific, Waltham, Massachusetts, United States) and the imaging system Fusion Fx7(Vilber, Collégien, France).

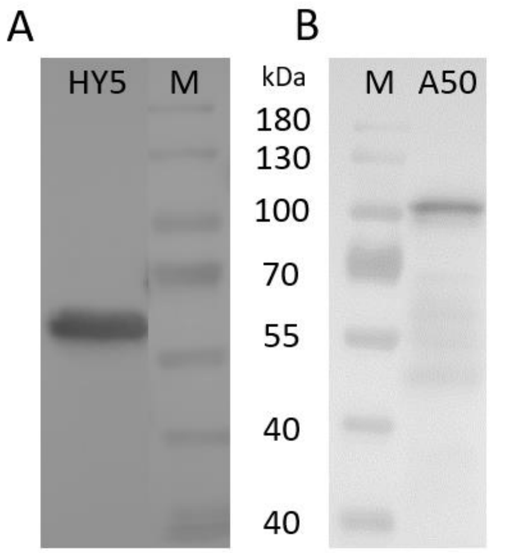


**Supplemental Figure 14: Western blot of Protein-Halo-Tag fusion**.

To confirm the expression of the HY5 (A) and ANAC050 (B) HaloTag® fusion proteins a SDS-Page with a 10% acrylamide gel followed by a western blot on a 0,22 µm Nitrocellulose membrane (Sartorius, Göttingen, Germany) was performed. The HY5 protein was expressed with TnT® SP6 High-Yield Wheat Germ Protein Expression System (Promega, Madison, Wisconsin, United States) using 2 µg plasmid DNA per 50 µL expression reaction (HY5). For size comparison the PageRuler™ (M) (Thermo Fisher Scientific, Waltham, Massachusetts, United States) was used. The Anti-HaloTag® Monoclonal Antibody (Promega, Madison, Wisconsin, United States) and an HRP conjugated anti-mouse antibody (abcam, Cambridge, United Kingdom) were used to detect the fusion protein. Detection was performed using Pierce™ ECL Western Blotting Substrate (Thermo Fisher Scientific, Waltham, Massachusetts, United States) and the imaging system Fusion Fx7(Vilber, Collégien, France). In the HY5 sample a band of approximately 60 kDa was observed. The ANAC050 sample shows a band just above 100 kDa. The expected sizes were 52 kDa for HY5 and 84 kDa for ANAC050. Both fusion proteins have a higher mass than expected. For HY5 this abnormal migration was already observed in previous studies[1].

**Table 1: EMSA binding conditions.** 10x Binding Buffer: 100 mM Tris, 500 mM KCl, 10 mM DTT, pH 7.5)

| | **Control** | **No competitor** | **Competitor** |
|---|---|---|---|
| **10X Binding Buffer** | 2 µl | 2 µl | 2 µl |
| **1M KCl** | 0.5 µl | 0.5 µl | 0.5 µl |
| **100 mM EDTA** | 0.5 µl | 0.5 µl | 0.5 µl |
| **200 mM MgCl2** | 0.5 µl | 0.5 µl | 0.5 µl |
| **Labelled probe** | 2 µl | 2 µl | 2 µl |
| **Competitor** | - | - | 4 µl |
| **Mut. Comp.** | - | - | - |
| **Wheat germ** | 8 µl (no plasmid) | 8 µl (320 ng plasmid) | 8 µl (320 ng plasmid) |
| **Nuclease free water** | 13.5 µl | 5.5 µl | 1.5 µl |
