## SupplementalMethods for "Local DNA shape is a general principle of transcription factor binding specificity in *Arabidopsis thaliana*"

**Pre-processing and training of the random forest regressor**

To perform data pre-processing and training of a random forest model an ampDAP-seq peak file (from http://neomorph.salk.edu/dap_web/pages/browse_table_aj.php) and the *Arabidopsis thaliana* genome (from arabidopsis.org) is necessary. Only peak files with FRiP value > 5% were considered and after motif extraction each peak file was filtered for peaks that appear in less then 66% of all ampDAP-seq available peak files, as those peaks were considered artifacts due to the ampDAP-seq procedure. Peak regions were extracted from the Arabidopsis genome using a custom python script, which expects one peak file (-p) and the corresponding genome (-g) as input. The resulting fasta file with genomic peak regions was used as input for the MEME-ChIP[1] (MEME-suite[2] v 5.0) tool with default parameters, so that the only given parameters were an output folder (-oc) and the peak regions fasta file (-dna). The sequence motif with highest e-value (--motif 1) from the resulting combined.meme file was searched in the *A. thaliana* genome using the FIMO[3] (MEME-suite[2] v 5.0) tool with a cut-off (--thresh) of 5e-4. To ensure that no matches were discarded the maximum number of stored matches (--max-stored-scores) were set to 1000000. The parameter --max-strand was set to 1 so that palindromic sequences would not match two times in the same locations and an output folder (--oc) was declared.

To allow a more accurate interpretation of binding affinities, the areas of motif matches were scanned for multiple motif occurrences using a custom python script. For this, each peak, which always has a length 200 base pairs, was tested for multiple FIMO matches. If more peaks had multiple motif occurrences than single motif occurrences, the corresponding transcription factor was considered for homodimeric binding events and vice versa. In that case, the random forest regressor was only trained on those peaks with multiple motif occurrences.

All genomic locations with sequence motif matches were translated into 13 DNA shape features using a publicly available query table[4], which was implemented into a custom Python script. Chloroplast and mitochondrial motif occurrences were discarded, as those sequences were not part of the in vitro experiment. The sequence window, for which the DNA shape was calculated, was set to 32 additional bases upstream and downstream from the sequence motif. Experimentally recorded signal values were normalized to range from 0 to 1000 using sklearns pre-processing module[5]. Since some shape features are mirrored for palindromic sequences, each matched sequence window on the minus strand was reverse complemented, so that the matrix of 3D shape values always correspond to the same direction.

DNA shape based training to learn protein binding affinities was performed with the RandomForestRegressor class from the sklearn module. The number of considered positions upstream and downstream of the sequence motif can be specified using the custom Python script by setting the -b parameter, which has a default value of 4. If the ratio of experimentally verified binding sites and genomic binding site occurrences was too high (> 1:5), this ratio was forced to be 1 to 5 by discarding random genomic positions. If this procedure would still yield more than 120,000 locations, the ratio was forced to be 1 to 3. This ratio between validated binding sites and binding site occurrences was always calculated for each transcription factor and used as sample weight for training the random forest regressor to prevent bias towards false negatives. The whole dataset was split into a train set (80%) and a validation set (20%) using the train_test_split function provided by the sklearn module, applying stratification to ensure even distributions of validated binding sites in the train and test set. The train set was used for 5-fold cross validation learning and the validation set was used for evaluation. To finetune the learning process, randomized hyperparameter grid search was performed for 75 iterations including the parameters "n-estimators" (ranging from 10 to 200), "max_features" (auto, square root or log2) and "max_depth" (ranging from 4 to 12). Additionally, 5-fold cross validation was performed, leading to a total number of 375 training procedures for each respective transcription factor. The mean squared error was used as loss. For evaluation purposes the precision recall curve function, which is also provided by the sklearn module, was applied on the validation data. After training the model, sequences of interest can be checked for putative binding affinity.
