## SupplementalFigure1 for "Local DNA shape is a general principle of transcription factor binding specificity in *Arabidopsis thaliana*"

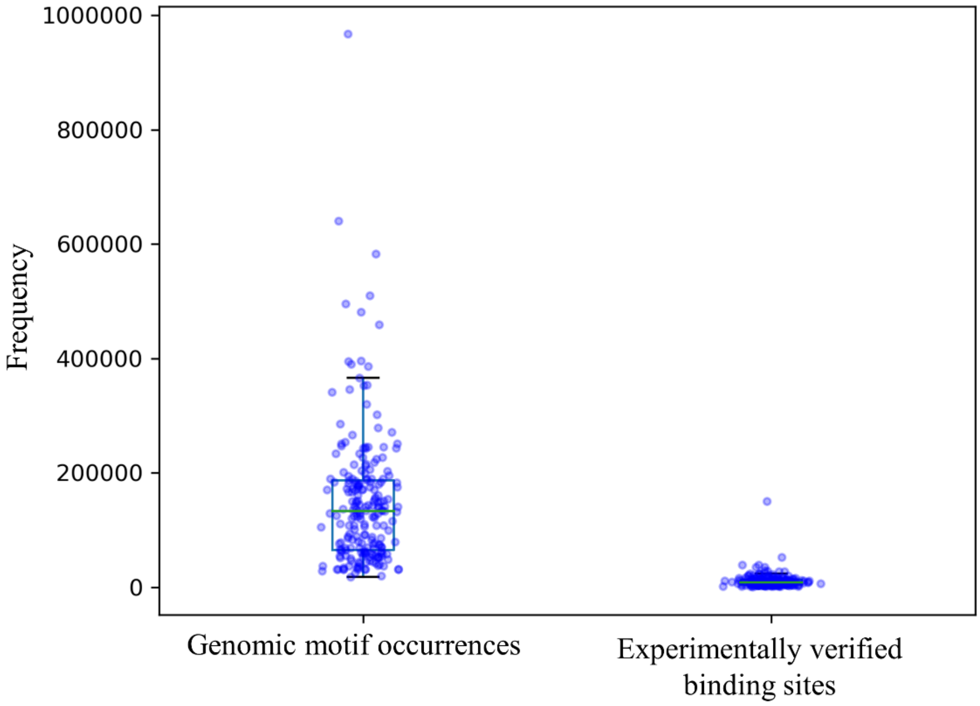


**Supplemental Figure 1: Comparison of motif occurrences and verified binding events.**

For the 216 tested transcription factors, actual binding events are on average 14-fold less frequent than binding sequence occurrences in the *A. thaliana* genome, for the respective transcription factor. The distribution on the left shows the number of individual motif occurrences for each transcription factor using the tool FIMO[1]. The distribution on the right shows the frequency of ampDAD-seq[2] verified binding events for each transcription factor.
