## SupplementalFigure2 for "Local DNA shape is a general principle of transcription factor binding specificity in *Arabidopsis thaliana*"

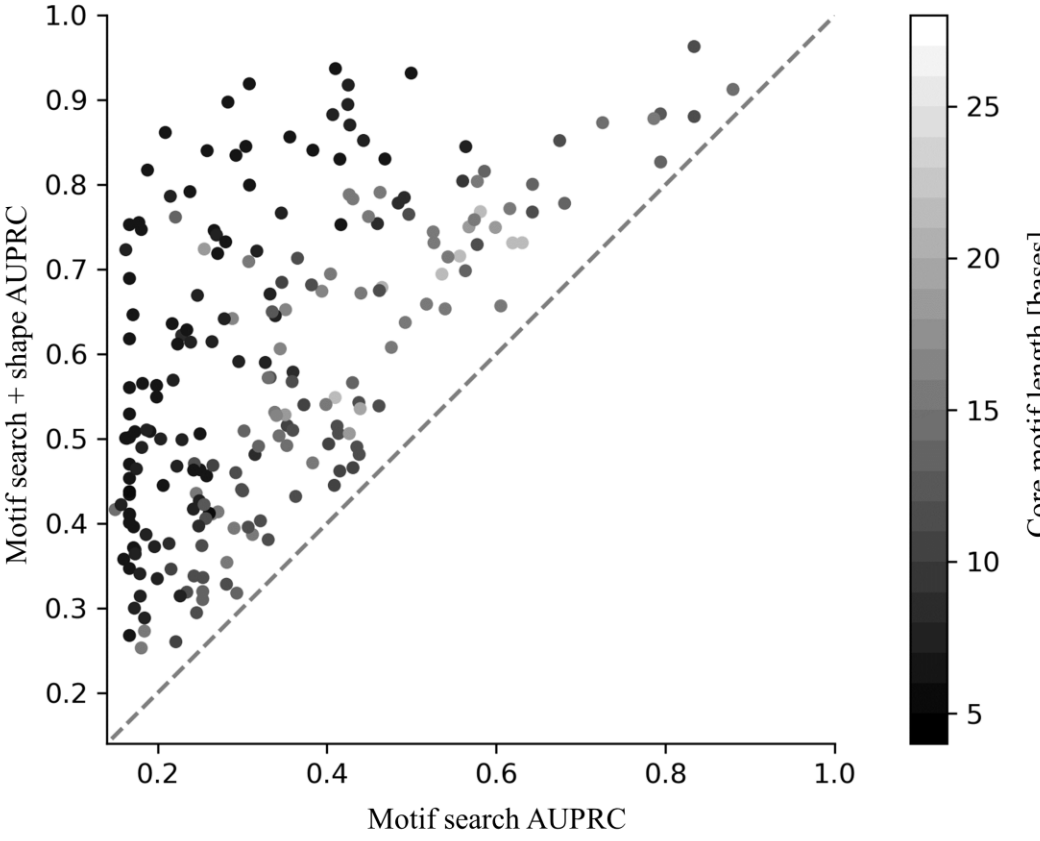


**Supplemental Figure 2: Comparison between sequence based motif search and adding a shape based classifier.**

The performance of identification of transcription factor binding sites improves for all tested transcription factors when adding shape information. Transcription factors with shorter sequence motifs, which are displayed as darker dots, tend to benefit most regarding the addition of DNA shape features.
