## SupplementalFigure3 for "Local DNA shape is a general principle of transcription factor binding specificity in *Arabidopsis thaliana*"

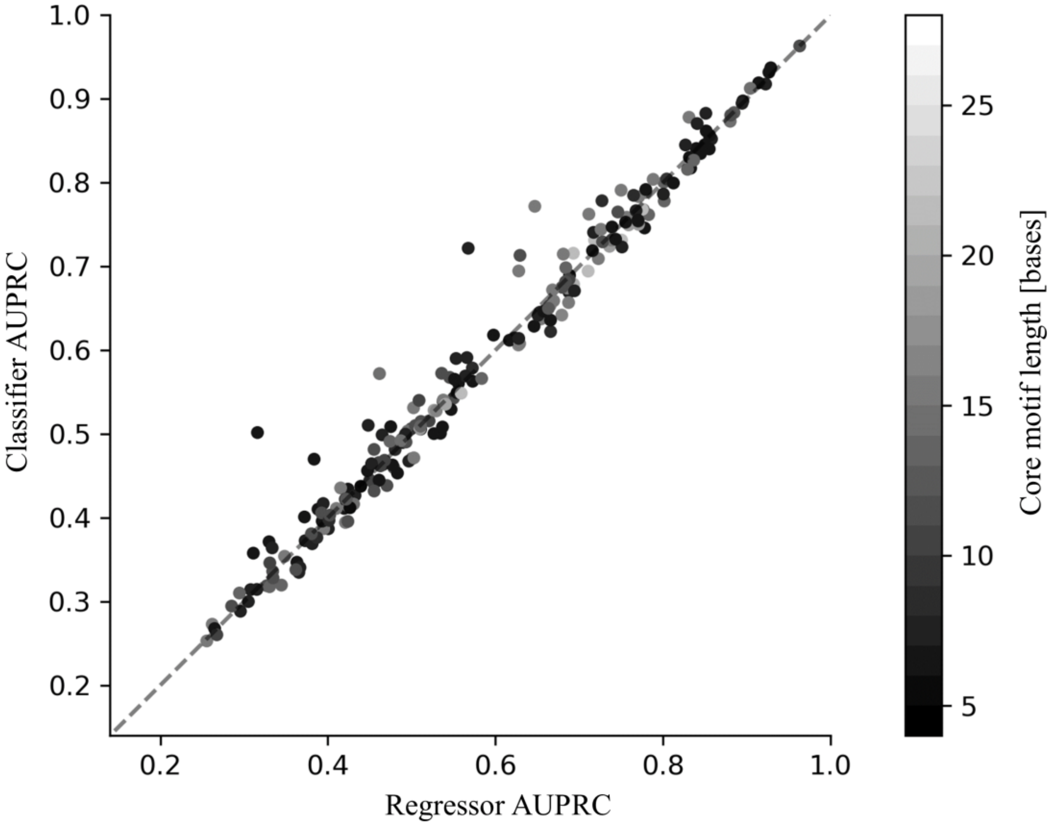


**Supplemental Figure 3: Comparison between random forest regressor and classifier.**

The binding site predictions of both approaches, represented by the AUPRC, are mostly similar. The classifier approach outperforms the regressor approach for specific transcription factors. However, the regression approach allows quantitative investigation of transcription factor binding events.
