## SupplementalFigure4 for "Local DNA shape is a general principle of transcription factor binding specificity in *Arabidopsis thaliana*"

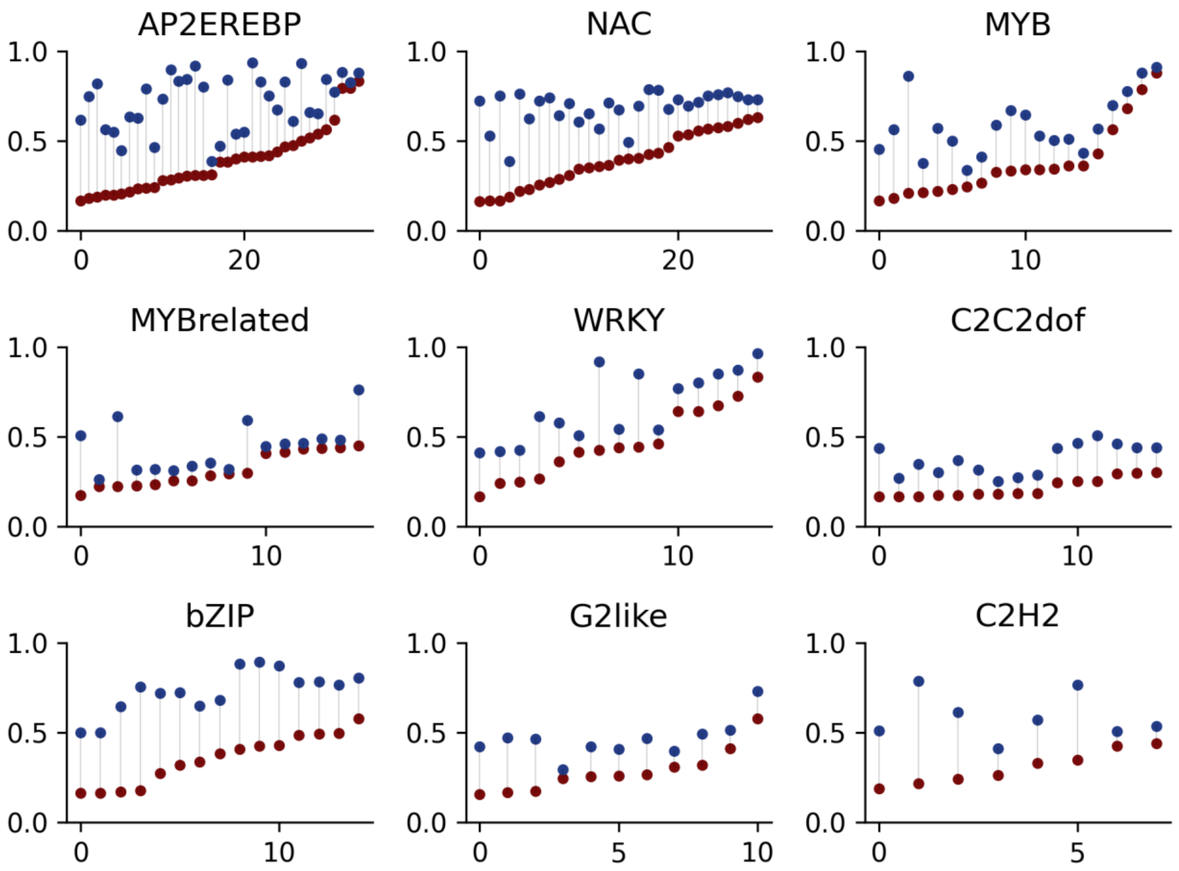


**Supplemental Figure 4: Individual binding prediction improvements regarding transcription factor protein families.**

Blue dots represent the area under the precision recall curve (AUPRC) using the regressor model which was trained on the DNA shape using DAP-seq data as ground truth, whereas red dots represent the AUPRC using solely the sequence motif derived from the ampDAP-seq peaks. For Members of the AP2EREBP, NAC and bZIP family, the binding prediction was consistently substantially improved. The improvement regarding the MYBrelated and C2C2dof family members was comparatively low.
