## SupplementalFigure5 for "Local DNA shape is a general principle of transcription factor binding specificity in *Arabidopsis thaliana*"

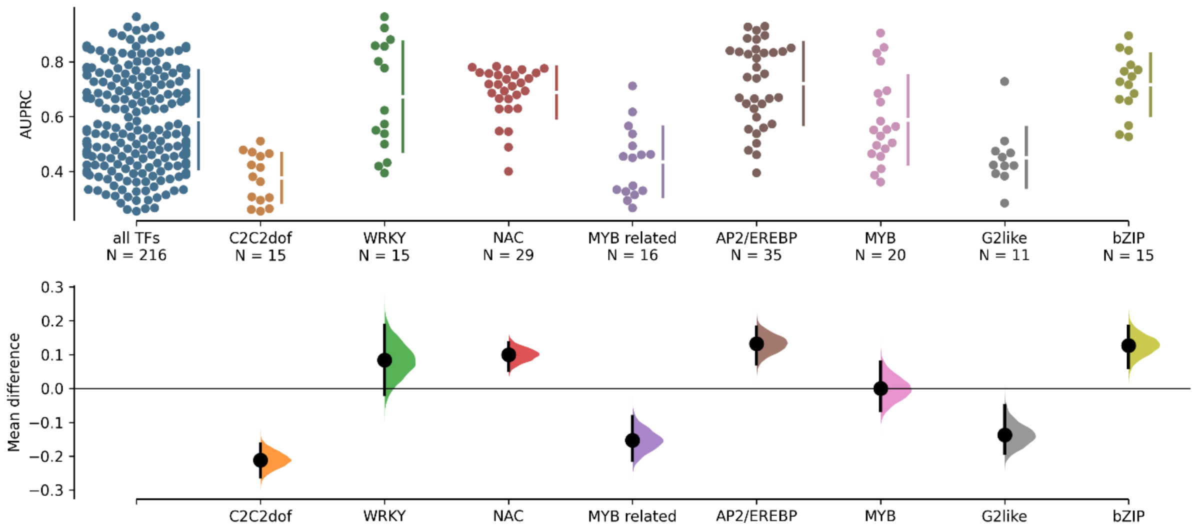


**Supplemental Figure 5: Comparative analysis of family specific DNA-binding prediction performance.**

The comparison is based on the area under the precision recall curve (AUPRC) using ampDAP-seq peaks as ground truth. Only families with available data of at least 10 members were investigated to ensure a robust analysis. For the families C2C2dof, MYB related and G2like a significantly ($p < 0.05$, Wilcoxon-Mann-Whitney-Test) worse performance of the random forest model regarding precision-recall relation was observed. For the transcription factor families NAC, AP2/EREBP and bZIP the performance was significantly ($p < 0.05$, Wilcoxon-Mann-Whitney-Test) better. The visualisation as well as the statistical tests were performed using the dabest package[1].

| Control | test | pvalue_mann_whitney |
|---|---|---|
| all TFs | C2C2dof | 0.000011 |
| all TFs | WRKY | 0.089488 |
| all TFs | NAC | 0.003943 |
| all TFs | MYB related | 0.000998 |
| all TFs | AP2/EREBP | 0.000065 |
| all TFs | MYB | 0.930433 |
| all TFs | G2like | 0.010457 |
| all TFs | bZIP | 0.006831 |
