## SupplementalFigure6 for "Local DNA shape is a general principle of transcription factor binding specificity in *Arabidopsis thaliana*"

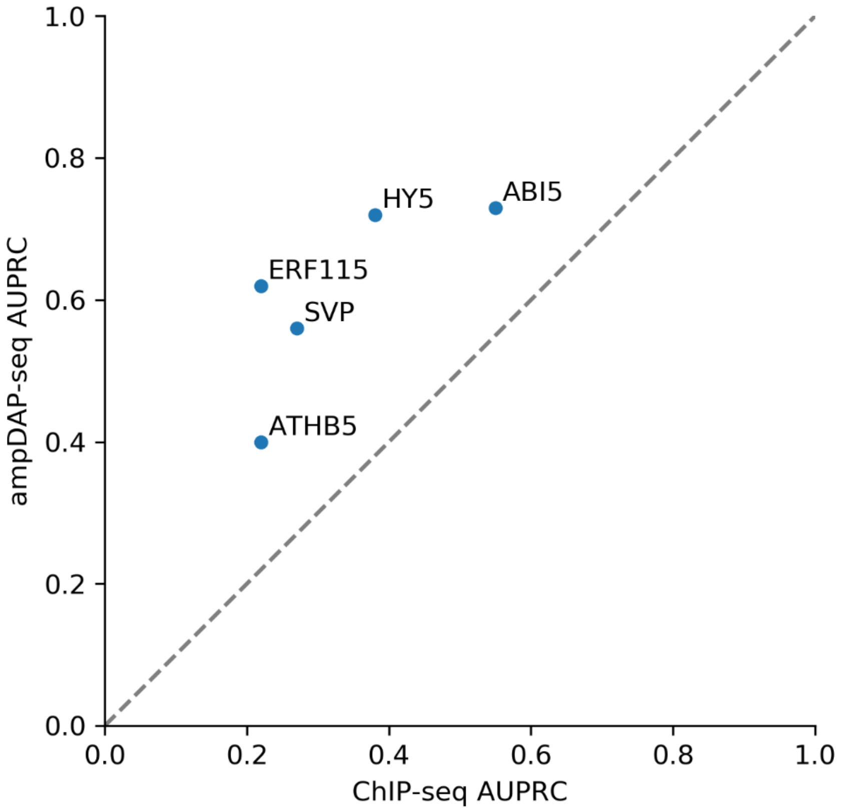


**Supplemental Figure 6: Comparison of ChIP-seq and ampDAP-seq data performance.**

For five transcription factors, which had data available for both experimental procedures[1–5], the performance of binding site inference was compared. For ChIP-seq the ChIP-seq data were used as ground truth for training and for ampDAP-seq the ampDAP-seq data were used as ground truth. The sequence motif was searched throughout the *Arabidopsis thaliana* genome to extract all putative binding sites. For each TF the random forest models trained on ampDAP-seq data had higher AUPRCs than models trained on ChIP-seq data.
