## SupplementalFigure7 for "Local DNA shape is a general principle of transcription factor binding specificity in *Arabidopsis thaliana*"

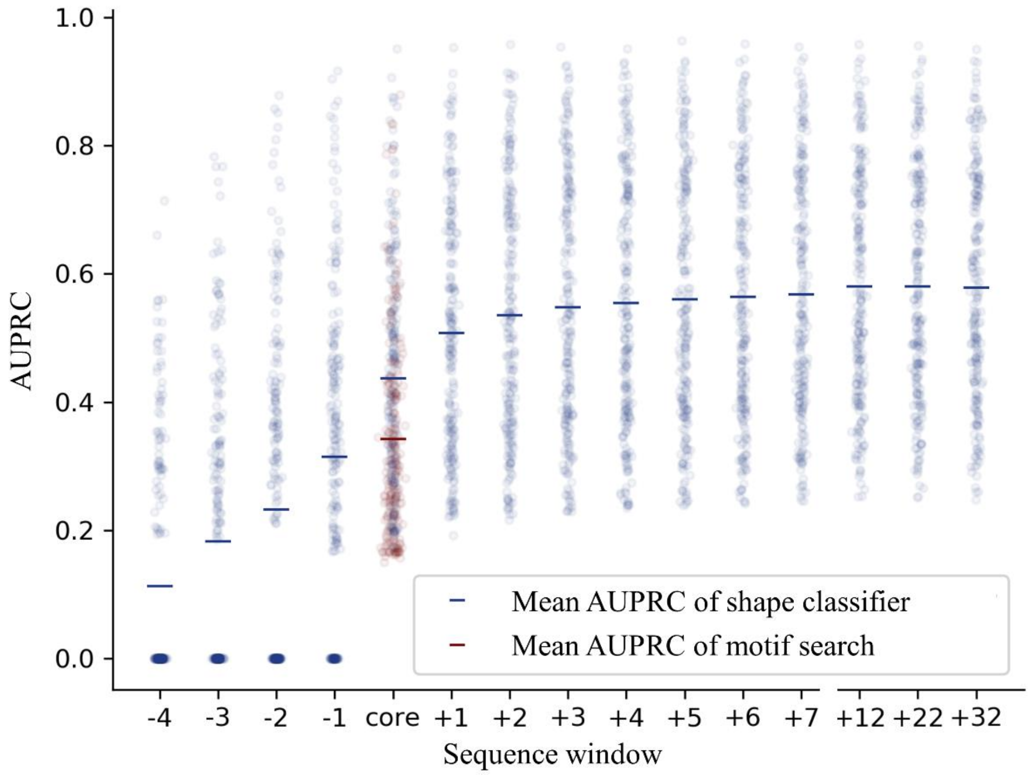


**Supplemental Figure 7: Binding site prediction performance for differing sequence windows.**

For each transcription factor the binding sequence window was varied up to 32 additional bases upstream and downstream from the core motif. The average AUPRC regarding the sequence search for the core motif is outperformed by calculating the DNA shape using only bases belonging to the core motif. Widening the sequence window with additional bases upstream and downstream from the core motif improves binding site prediction to an average AUPRC of approximately 0.6. The spread of prediction performance varies widely and is dependent on each transcription factor.
